# Growing out of the fins: implications of isometric and allometric scaling of morphology relative to increasing mass in blue sharks (*Prionace glauca*)

**DOI:** 10.1101/2023.12.21.572684

**Authors:** Scott G. Seamone, Phillip C. Sternes, Theresa M. McCaffrey, Natalie K. Tsao, Douglas A. Syme

## Abstract

Disproportional changes (i.e. allometry) in shark morphology have been attributed to shifts in function associated with niche shifts in life history, such as in habitat and diet. Photographs of blue sharks (*Prionace glauca,* 26-145 kg) were used to analyze changes in parameters of body and fin morphology with increasing mass that are fundamental to swimming and feeding. We hypothesized that blue sharks would demonstrate proportional changes (i.e. isometry) in morphology with increasing mass because they do not undergo profound changes in prey and habitat type, but as a result, we predicted that blue sharks would grow into bodies with greater turning inertias and smaller frontal and surface areas, in addition to smaller spans and areas of the fins relative to mass. Many aspects of morphology increased with isometry. However, blue sharks demonstrated negative allometry in body density, whereas surface area, volume and roll inertia of the body, area, span and aspect ratio of both dorsal fins, span and aspect ratio of the ventral caudal fin, and span, length and area of the mouth increased with positive allometry. The dataset was divided in half based on mass to form two groups: smaller and larger sharks. Besides area of both dorsal fins, relative to mass, larger sharks had bodies with significantly greater turning inertia and smaller frontal and surface areas, in addition to fins with smaller spans and areas, compared to smaller sharks. Hence, isometric scaling does not necessarily imply functional similarity, and allometric scaling may sometimes be critical in maintaining, rather than shifting, function relative to mass. Both allometric and isometric changes in blue sharks are predicted to promote reduced costs of transport in migration, but conversely, decreased unsteady performance, such as in escape responses. These changes are likely beneficial for larger sharks that probably experience reductions in predation pressure.

## INTRODUCTION

Exploring how morphology changes with increasing body size (e.g. mass, length, height) in organisms is the study of scaling (Gayon, 2000; Huxley and Tessier, 1936; Schmidt-Nielsen, 1975). Proportional relationships between the scaling of morphology and size are referred to as isometric growth or isometry, which was derived from the Greek words ‘*isos’* and ‘*metron*’, meaning equal and measure, respectively (Gayon, 2000; Huxley and Tessier, 1936; Schmidt-Nielsen, 1975). In contrast, disproportional changes in morphology with size are referred to as allometric growth or allometry (Gayon, 2000; Huxley and Tessier, 1936; Schmidt-Nielsen, 1975). This term was originated by Huxley and Tessier (1936) to avoid confusion in the nomenclature of relative growth, and was derived from the Greek ‘*alloios*’, which means different (Gayon, 2000; Schmidt-Nielsen, 1975). Ever since Galileo Galilei (1637) revealed disproportionate growth relative to increasing mass in the skeleton of large mammals such as the elephant, scaling has become an informative field of study for biologists, providing insight into both intraspecific and interspecific relationships including the impacts of size on metabolism, feeding and locomotion in terrestrial and aquatic organisms (Birn-Jeffery and Higham, 2014; Cook, 1996; Cullen and Marshall, 2019; Dial et al., 2008; Domenici, 2001; Enquist et al., 2003; Gould, 1966; Higham et al., 2018; Killen et al., 2007; Norin and Gamperl, 2018; Vogel, 2005). Of relevance to the present research, scaling studies can have utility for advancing our understanding of the relationships between form, function, behaviour and ecology in species that can be difficult to study in their natural habitat, such as large sharks that swim in oceanic (i.e. deep bodied) waters beyond the continental shelf (Ahnelt et al., 2020; Gayford et al., 2023a; Sternes and Higham, 2022; Yun and Watanabe, 2023).

There has been a recent growth of interest in scaling relationships in sharks (Ahnelt et al., 2020; Bellodi et al., 2023; Fu et al., 2016; Gayford et al., 2023a; Gayford et al., 2023b; Gleiss et al., 2017; Irschick and Hammerschlag, 2015; Irschick et al., 2017; Lingham-Soliar, 2005; Reiss and Bonnan, 2010; Sternes and Higham, 2022; Yun and Watanabe, 2023). The liver size in sharks can increase with positive allometry, both interspecifically and intraspecifically (Bone and Roberts, 1969; Gleiss et al., 2017; Lingham-Soliar, 2005). Sharks are negatively buoyant, and an allometric increase in the liver mass, which has a high fat content and thus contributes buoyancy, suggests larger sharks might become more neutrally buoyant with increasing size, which is anticipated to have impacts on the energetics of swimming (Gleiss et al., 2017; Iosilevskii and Papastamatiou, 2016). Many aspects of body and fin morphology tend to be isometric relative to increasing length of the shark (Irschick and Hammerschlag, 2015; Irschick et al., 2017). However, in coastal species that show habitat and diet shifts throughout their life history, allometric changes in head and fin shape with increasing length have been revealed; this has been attributed to potential differences among adult and juvenile sharks in locomotor and predation strategies (Fu et al., 2016; Gayford et al., 2023a; Irschick and Hammerschlag, 2015; Irschick et al., 2017; Lingham-Soliar, 2005; Sternes and Higham, 2022). Gayford et al. (2023a) recently proposed the ‘allometric niche shift’ hypothesis which suggests selective pressures related to ontogenetic changes in diet and habitat may represent important drivers of allometric scaling in the morphology of sharks.

Belonging to the Order Carcharhiniformes, blue sharks (*Prionace glauca*, Linnaeus 1758) are a highly studied species commonly found in oceanic waters from temperate to tropical temperatures in the Atlantic, Pacific and Indian Ocean (Ebert et al., 2021). A large size range is noted in these sharks, such that pups can grow from about 35cm in length and 1kg in mass into adults larger than 380cm and 200kg (Ebert et al., 2021; International Game Fish Association, 2002; Kohler et al., 1992). Stable isotope studies in blue sharks demonstrate differences between populations whereby trophic level and foraging location may vary with size (Estrada et al., 2003; Hernandez-Aguilar et al., 2016; Kiszka et al., 2015; Li et al., 2016; Macneil et al., 2005; Preti et al., 2012; Rabehagasoa et al., 2012; Vidal et al., 2023). However, blue sharks seem to be generalist predators whereby the range of prey type appears to be relatively consistent throughout their life history (Bazzi et al., 2021; Henderson et al., 2001; McCord and Campana, 2003; Young et al., 2010). Rather than profound changes with growth in predatory behaviour and prey type as observed in tiger and white sharks (Fu et al., 2016; James et al., 2006; Lingham-Soliar, 2005; Lowe et al., 1996), trophic level shifts in blue sharks have been attributed to availability of prey in different foraging locations and larger individuals having access to larger prey (Hernandez-Aguilar et al., 2016; Rabehagasoa et al., 2012). With regards to habitat, blue sharks never cease swimming in deep bodied and wide-open oceanic waters throughout their life history (Campana et al., 2011; Queiroz et al., 2005; Stevens, 1976; Vandeperre et al., 2014). Although larger individuals exhibit a wider migratory range than smaller juveniles that appear to remain near nursery grounds, the distance travelled as a function of time is apparently similar across life history (Vandeperre et al., 2014). Furthermore, blue sharks do not seem to undergo major habitat shifts compared to species like hammerheads, which migrate from shallow mangrove environments to oceanic waters offshore (Clarke, 1971; Colombo Estupiñán-Montaño et al., 2021; Duncan and Holland, 2006; Hoyos-Padilla et al., 2014; Sternes and Higham, 2022). Hence, the physical environment experienced by blue sharks in oceanic waters is assumed to be relatively consistent between juveniles and adults.

Due to the lack of substantial changes in the type of prey and physical environment that blue sharks seem to experience throughout their life history, we hypothesized that these sharks would exhibit isometric growth in body form and fin shape with increasing mass. However, according to geometric scaling laws, linear and area dimensions intrinsically decrease relative to increasing mass with isometric growth (Galileo, 1914; Schmidt-Nielsen, 1975). Accordingly, we further predicted that relative to mass, isometric growth would lead to significantly smaller planform spans and areas of the fins, smaller frontal and surface areas of the body, and greater turning inertias of the body in larger versus smaller sharks. As such, one of our objectives was to explore the idea that isometric scaling is not necessarily associated with functional similarity in sharks that exhibit an extensive range in body size, and to infer the prospective implications of isometry on the energetic cost of transport in migration (Di Santo et al., 2017; Gamperl et al., 2002; Lauder and Di Santo, 2015; Sepulveda et al., 2007; Watanabe et al., 2019) and the performance of unsteady burst swimming such as in escape responses in sharks (Domenici, 2001; Domenici and Blake, 1997; Domenici et al., 2004; Seamone et al., 2014). To test the hypothesis and ideas, we used photographic images of blue sharks to analyze parameters of body and fin shape that are fundamental to swimming and feeding, and how they change relative to mass as these sharks grow.

## MATERIALS AND METHODS

### Data collection

Data collection was approved by the Animal Care Committee at the University of Calgary under protocol AC15-0065, following guidelines of the Canadian Council on Animal Care. Photographs were obtained of thirty-six blue sharks (*Prionace glauca,* Linnaeus 1758), freshly caught by fisherman with hook and line from the Atlantic Ocean off the coast of Nova Scotia, Canada, during shark derbies sanctioned by Fisheries and Oceans Canada. Sharks were sacrificed immediately after being caught by derby participants, according to Fisheries and Oceans Canada guidelines, and placed on ice while they were brought to the weighing docks. Total mass was measured for each individual, whereby sharks were hung from a scale vertically from their caudal peduncle. Sex was determined by the presence of claspers, and data from a total of 16 males and 20 females were recorded. Sharks were then placed with the ventral surface on the ground, and photographs were taken using a Panasonic Lumix DMC-FZ1000 digital camera (image resolution 20.10 megapixels; Panasonic Corporation, Kadoma, Japan), with a scale (35.56 cm) placed square to the camera in each image for calibration. Photographs included a lateral view of the entire shark, whereby the shark spanned the entire field of view of the camera. Then from anterior to posterior, three separate but overlapping photographs of the dorsal surface of the shark were taken, whereby the camera was approximately 1.5m above the specimen. Analyses included images from dorsal and caudal fins held in position during photographs. The pelvic and anal fins were not accessible for these photographs.

### Data analysis

Images were analyzed using ImageJ (bundled with 64-bit Java 1.8.0_112, Wayne Rasband Developers 1997) and the results then imported into Excel spreadsheets to quantify morphological parameters relevant to shark locomotion and feeding, as described below. Locations of the different measurements used are shown in Figure 1.

**Figure 1.**
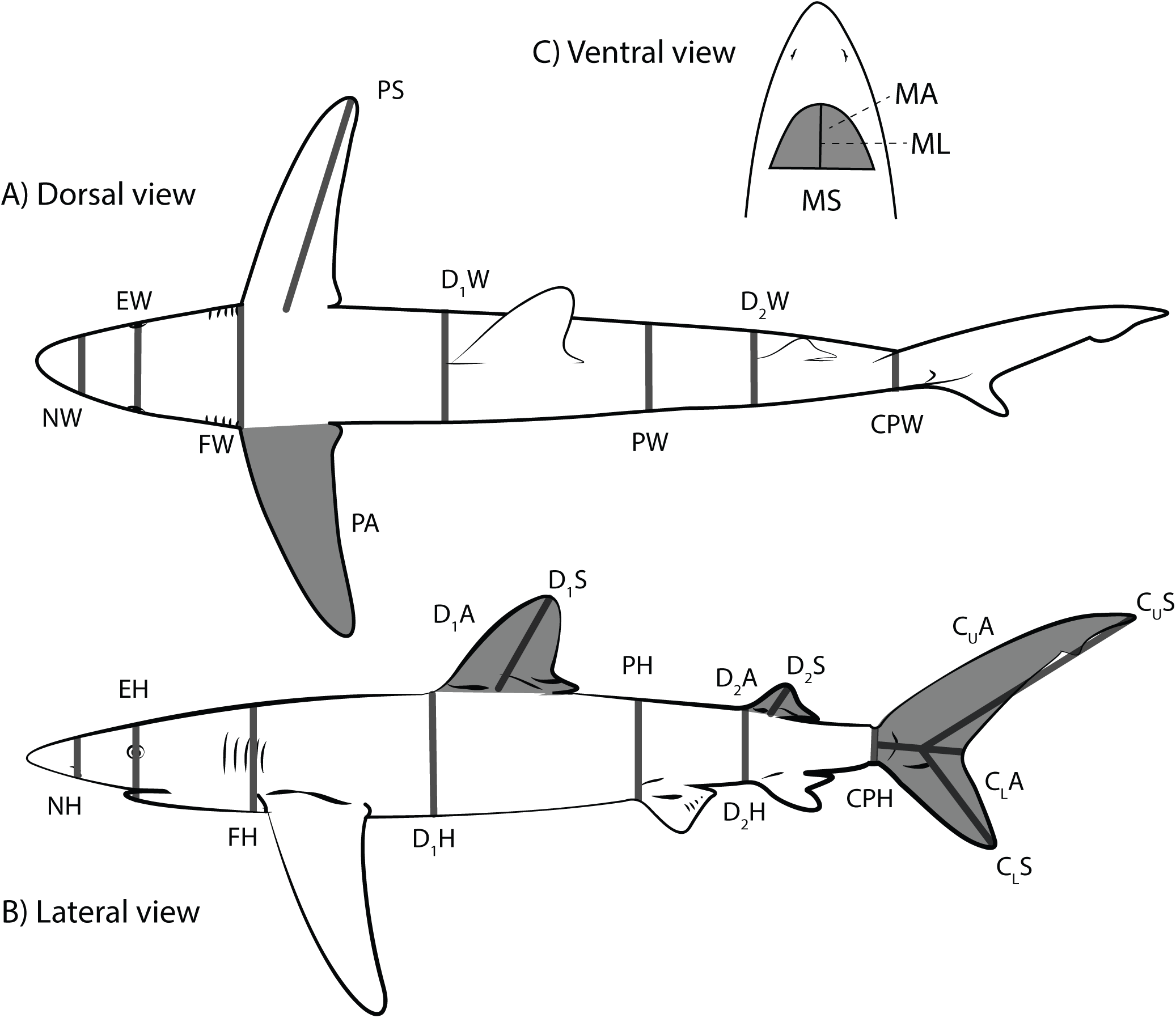
Morphological measurements of the blue shark (H = height, W = width, S = span, C_U_ = upper caudal, and C_L_ = lower caudal, CP = caudal peduncle, D_1_ = first dorsal, D_2_ = second dorsal, E = eye, F = frontal, N = nostril, P = pelvic, M = mouth).

#### Body volume and density

Density was calculated by dividing the total mass by the volume of the shark. Volume of the shark was calculated using the equation for the volume of a cone (⅓πR^2^L) from the nostril to the snout, and then for the volume of a partial cone (⅓πL(R_1_^2^+ R_1_R_2_+ R_2_^2^)) for each segment of the shark from the nostril to the caudal peduncle, where R is the average radius of the base of the cone determined from the height and width measurements at a give section of the shark. Hence, R_1_ and R_2_ indicate the average radius for each base of the partial cone for each segment. The volume for each segment was then summed for the total volume of the body.

#### Fineness ratio and area of the body

Fineness ratio, used as an index to quantify streamlining of the body of the shark, was measured as the standard length of the body divided by the maximum width of the body (i.e. the width anterior to the pectoral fins). While swimming, drag is a force acting opposite to the relative motion of the shark with respect to the surrounding water, where less drag is associated with more energetically economic swimming and the capacity to reach higher sustained swimming speeds (Blake, 1983; Videler, 1993; Webb, 1975). Pressure drag appears to be the dominant form of drag experienced by large aquatic vertebrates and occurs due to distortion in the flow of water around the body (Webb, 1975). Pressure drag is proportional to the maximum area of the frontal profile of the shark (Webb, 1975). The frontal area was measured as the transverse area of an assumed ellipse (πR^2^), where R was measured as the average radius of the height (from the lateral images) and width (from the dorsal images) of the body at the leading edge of the pectoral fins. Frictional drag is another type of drag that occurs due to the friction between the water and the skin of the shark, and is proportional to the surface area of the shark (Blake, 1983; Videler, 1993; Webb, 1975). Surface area was measured as the surface area of a cylinder (2πRL+2πR^2^), where R was calculated as the average radius of an assumed cylinder (√(Vπ^-1^L^-1^)) using the body volume (V) measurements described above and the standard length (L) of the shark.

#### Moment of inertia

Moment of inertia, i.e., turning inertia in a swimming fish, describes a quantity expressing a body’s tendency to resist angular acceleration (i.e. roll, pitch, and yaw), whereby a greater moment of inertia is equivalent to greater inherent stability (less tendency to deflect and turn). Assuming the location of the centre of mass does not change and the shape of the body is relatively similar as sharks grow, the product MR^2^ provides an appropriate index for the moment of inertia of roll, pitch and yaw, where M equals the mass of the shark and R is the perpendicular distance of the applied torque to the axis of rotation (i.e. the moment arm). For roll, R was measured as the average radius of an assumed cylinder (√(Vπ^-1^L^-1^)) using the body volume (V) measurements described above and the length (L) of the shark. For pitch and yaw, R corresponded to the standard length of the animal, and hence, indices for moment of inertia for yaw and pitch are the same.

#### Fins

The fins of sharks generate forces (i.e. lift and drag) to promote stability and maneuverability when swimming (Hoffmann and Porter, 2019; Maia and Wilga, 2013a; Maia and Wilga, 2013b; Standen, 2005; Standen, 2008; Wilga and Lauder, 2000). Fin span, which also is an index for the moment arm of the fin, was measured as the linear distance from the mid point of the fin base to the tip of the fin. Planform area, which also provides an index of the magnitude of force along the fins (Blake, 1983; Videler, 1993; Webb, 1975), was measured by tracing the perimeter of both the upper and lower lobes of the caudal fin and the anterior and posterior dorsal fins from the lateral images, while the pectoral fins were measured from the dorsal images. For both span and area, left and right pectoral fins were measured for 10 sharks to confirm there was no significant asymmetry, and then one fin was measured for each successive shark.

Aspect ratio was measured as span^2^/planform area of the fins (Blake, 1983; Videler, 1993; Webb, 1975). The aspect ratio of these surfaces is related to their lift:drag ratio, whereby surfaces with a higher aspect ratio have the capacity to remain stable at higher speeds, while lower aspect ratios promote greater maneuverability (i.e. procured acceleration) (Blake, 1983; Lighthill, 1969; Webb, 1975; Webb, 1984).

#### Head and mouth

Changes in the shape of the head and mouth might imply differences in bite force and prey handling with growth (Fu et al., 2016). Dorsal view images were used to measure area and span of the head. Span of the head was measured as the maximum linear distance anterior to the pectoral girdle. Head area was measured by tracing the perimeter of the head anterior to the pectoral girdle. Aspect ratio of the head was used to determine if the basic shape of the head was changing and was measured as span^2^/area of the head. Ventral view images were used to measure span, length and area of the jawline of the mouth. The jawline when closed formed a semicircle-like shape. The span of the mouth was measured as the maximum linear distance from one side of the jawline to the other. Length of the mouth was measured as the linear distance from the line crossing the width of the jawline to the arch of the semicircle. The area of the mouth was measured by tracing the area encompassed by the semicircle and width of the lower jawline.

#### Scaling and statistics

All statistics were performed in R software (R Development Core Team 2008). To test the hypothesis that the shape of blue sharks scaled with isometry (i.e. geometrically similar) with increasing mass, morphological parameters were scaled according to the power-law function *y*=*aM^b^*, where *M* is body mass (in kilograms) and *b* is the scaling exponent. All data were Log_10_ transformed prior to analyses and then related to log_10_*M* using a least-squares linear regression model. Scaling equations then took the form: log*y*=log*a*+*b*log*M*, whereby the scaling coefficient, b, is the slope of the relationship between log*M* and the log of the variable under study.

Isometric organisms are expected to have linear dimensions that scale with *M^1^*^/3^, measures of area proportional to *M*^2/3^, measures of mass and volume proportional to *M*^1^, and measures of density proportional to *M*^0^. Aspect ratio (i.e. span^2^/area) is unitless and is thus expected to scale to M^0^ for isometry, while moment of inertia (i.e. MR^2^) is expected to scale to M^5/3^ for isometry. To compare the scaling exponents obtained to those expected based on isometry, the 95% confidence interval of the exponent was calculated. If the 95% confidence interval for the measured exponent encompassed the value expected for isometry the relationship was considered isometric, but if the confidence interval was less than the expected value the relationship was considered negative allometry, or positive allometry if the confidence interval was greater than the expected value. Males and females were pooled together for regression testing, because the females did not cover a great enough mass range to be reliably tested alone. To ensure the pooling of sexes did not notably skew the results, the males were also tested alone, and results were compared with the pooled data. Differences in scaling between the pooled vs male-only data sets was observed in the mouth length and area. Pooled data scaled with a positively allometric slope for both variables with significant differences from isometry. Male only data scaled with a positively allometric slope, but the 95% confidence interval was not significantly different than the slope for isometry (see Results). Because the lack of difference from isometry was slight, we proceeded with interpreting the functional implications of the results for the pooled data.

To test the prediction that scaling would lead to significant differences in the planform area and span of the fins, the frontal and surface areas of the body, and the yaw, pitch and roll moment of inertias of the body relative to mass in larger versus smaller sharks, the dataset was divided in half based on the mass of the sharks to form two groups: small sharks and large sharks. The magnitude of the span and area of the fins, surface and frontal areas of the body, and moment inertias of the body were then normalized by the mass of the shark. Afterwards, two sample t-tests were used to compare differences in the means for the variables in larger versus smaller sharks.

## RESULTS

Mass of male sharks ranged from 26.7 to 144.8 kg (mean 90.2 ± 8.36 SEM), and standard length ranged from 162.2 to 277.1 cm (mean 228.6 ± 8.78 SEM), while mass of female sharks ranged from 26.3 to 46.8 kg (mean 34.7 ± 1.52 SEM) and standard length ranged from 138.2 to 170.9 cm (mean 156.6 ± 2.76 SEM). For the body, standard length, fineness ratio, frontal area, and pitch and yaw inertia increased with isometry (Table 1), whereas surface area, volume, and roll inertia increased with positive allometry (Figure 2, Table 1), and density increased with negative allometry (Figure 2, Table 1). For the caudal fin, dorsal span, dorsal area, dorsal aspect ratio, and ventral area increased with isometry (Table 1), whereas ventral span and ventral aspect ratio increased with positive allometry (Figure 3, Table 1). For both dorsal fins, span, area, and aspect ratio increased with positive allometry (Figure 4 and 5, Table 1). For the pectoral fins, span, area, and aspect ratio increased with isometry (Table 1). For the head, span, area, and aspect ratio increased with isometry (Table 1). For the mouth, span, length, and area increased with positive allometry (Figure 6, Table 1), whereas aspect ratio increased with isometry (Table 1). The slope of male only data for the length of the mouth increased with positive allometry but was sightly not significantly different than isometry (p-value < 0.001, R^2^ = 0.74, slope ± CI = 0.43 ± 0.10). The slope of male only data for the area of the mouth increased with positive allometry but was slightly not significantly different than isometry (p-value < 0.001, R^2^ = 0.81, slope ± CI = 0.80 ± 0.15). Normalized by mass, the span and area of the fins except for the area of both dorsal fins, in addition to the frontal area and surface area of the body was smaller, whereas roll, pitch and yaw inertia of the body was greater in the grouped larger versus smaller sharks (Table 2). The relative change in variables that scaled with allometry as opposed to isometry are displayed as percentages in Table 3.

**Figure 2.**
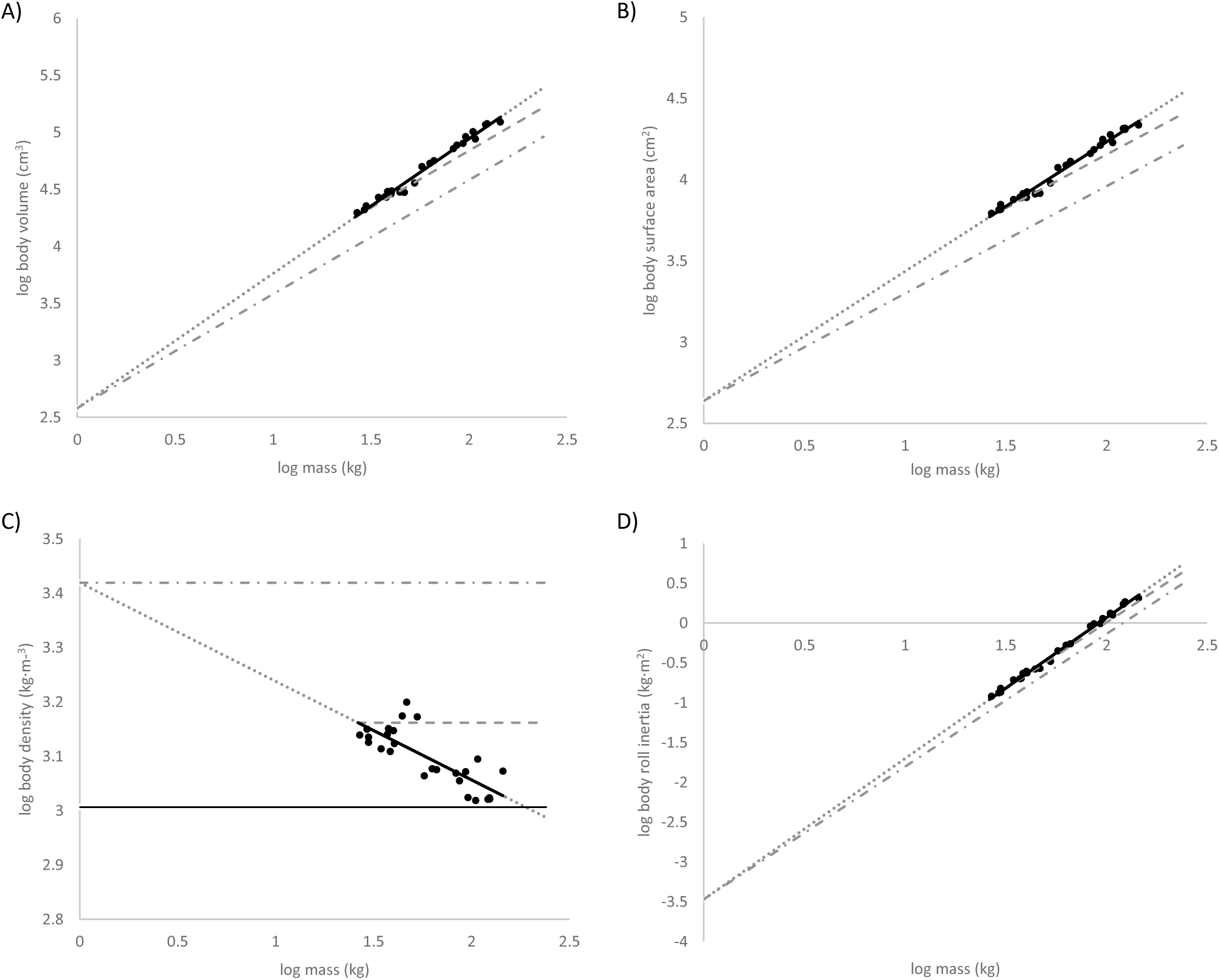
Allometric scaling of the body, represented by the black dots and solid line. Dotted line represents the allometric scaling extended across the size range observed in blue sharks (1kg-240kg). Dashed line represents isometric scaling initiated from the smallest shark observed in the dataset of this study (26kg). Dashed and dotted line represents isometric scaling initiated from the size of a blue shark pup (1 kg). A) Volume (allometric slope = 1.18, R^2^ = 0.98; isometric slope = 0.33). B) Surface area (allometric slope = 0.80, R^2^ = 0.98; isometric slope = 0.66). C) Density (allometric slope = −0.18, R^2^ = 0.61; isometric slope = 0). D) Roll inertia (allometric slope = 1.76, R^2^ = 0.99; isometric slope =1.66).

**Figure 3.**
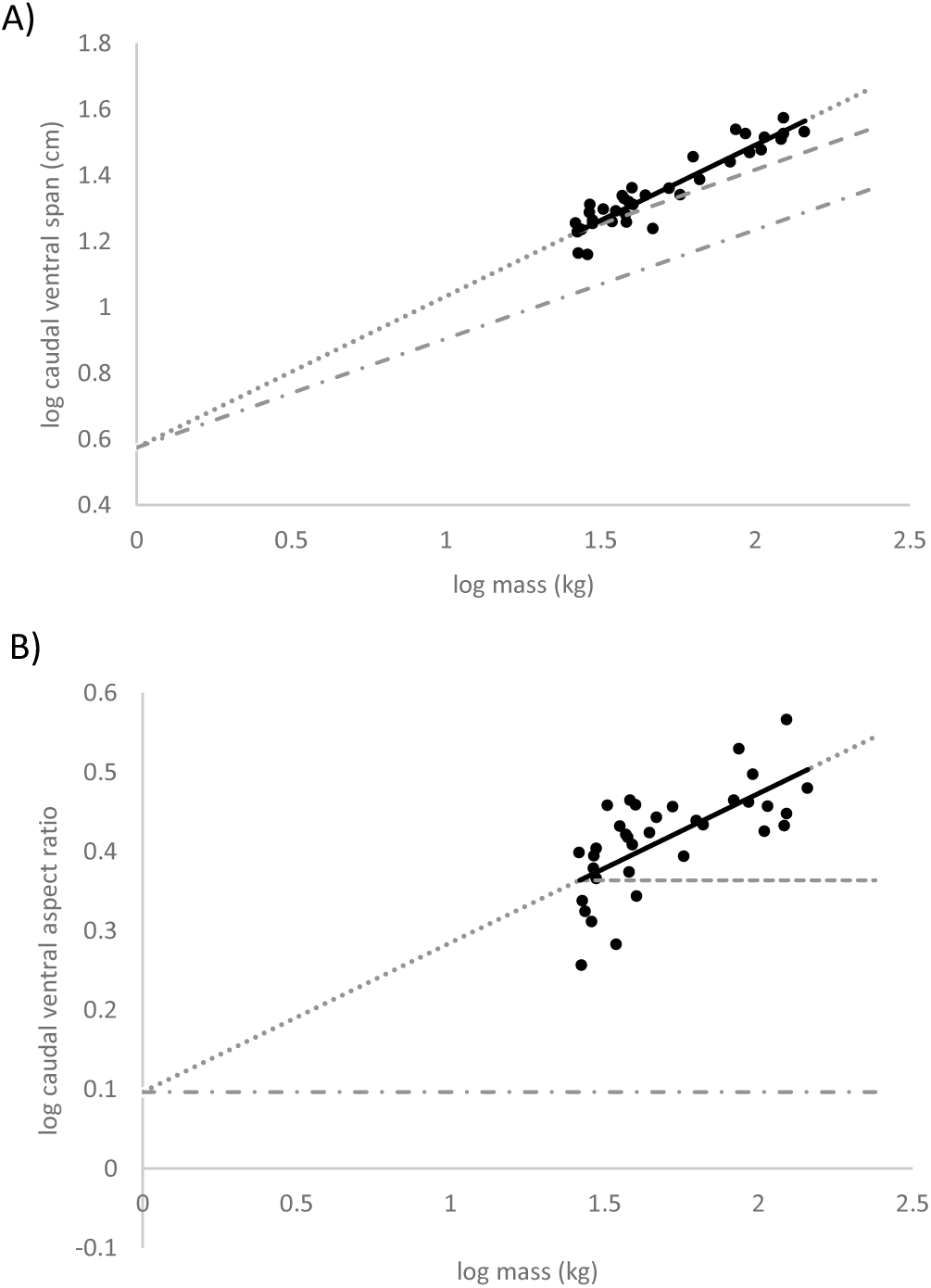
Allometric scaling of the ventral caudal fin, represented by the black dots and solid line. Dotted line represents the allometric scaling extended across the size range observed in blue sharks (1kg-240kg). Dashed line represents isometric scaling initiated from the smallest shark observed in the dataset of this study (26kg). Dashed and dotted line represents isometric scaling initiated from the size of a blue shark pup (1 kg). A) Fin span (allometric slope = 0.46, R^2^ = 0.87; isometric slope = 0.33). B) Fin aspect ratio (allometric slope = 0.19, R^2^ = 0.45; isometric slope = 0).

**Figure 4.**
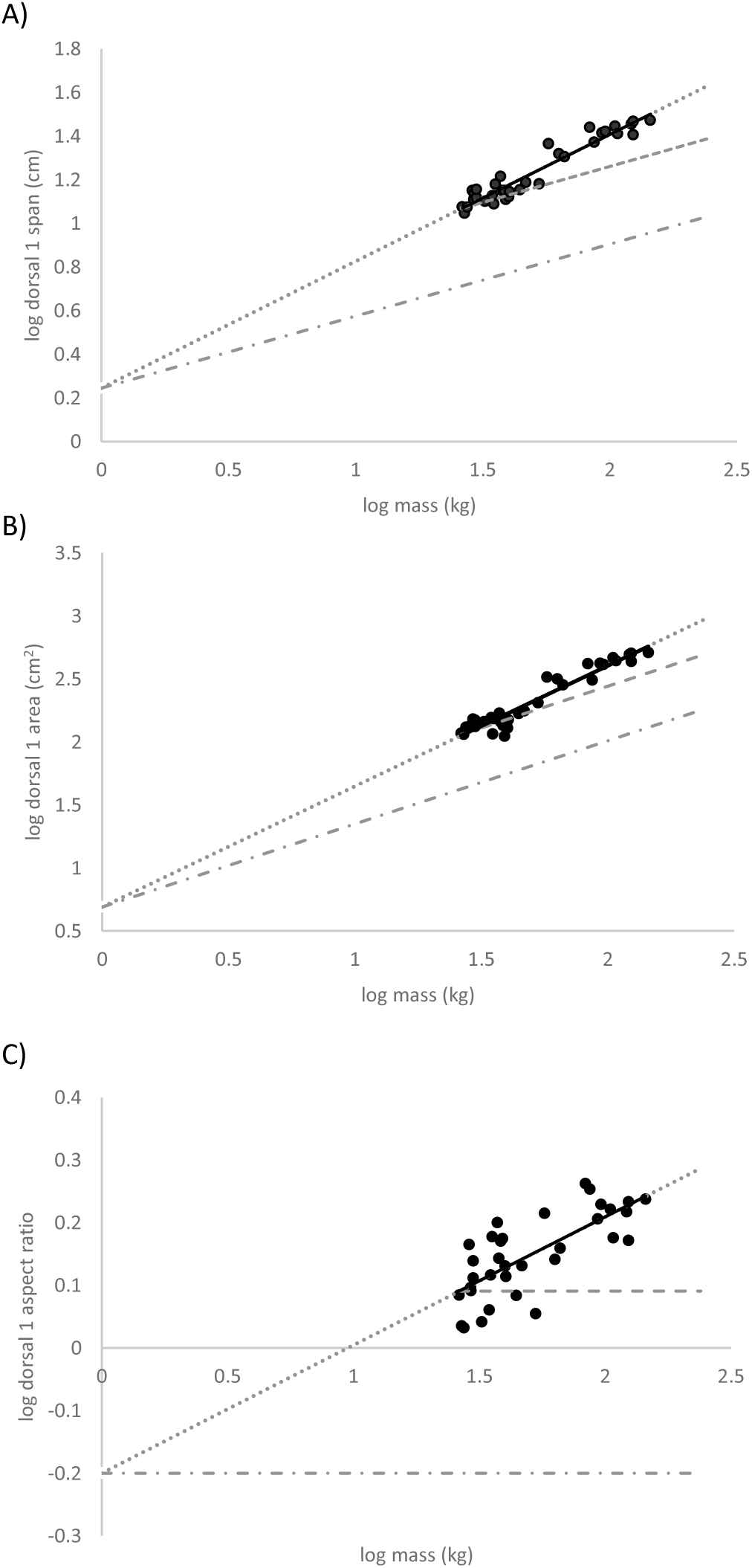
Allometric scaling of the first dorsal fin, represented by the black dots and solid line. Dotted line represents the allometric scaling extended across the size range observed in blue sharks (1kg-240kg). Dashed line represents isometric scaling initiated from the smallest shark observed in the dataset of this study (26kg). Dashed and dotted line represents isometric scaling initiated from the size of a blue shark pup (1 kg). A) Fin span (allometric slope = 0.58, R^2^ = 0.92; isometric slope = 0.33). B) Fin area (allometric slope = 0.96, R^2^ = 0.92; isometric slope = 0.66). C) Fin aspect ratio (allometric slope = 0.21, R^2^ = 0.53; isometric slope = 0).

**Figure 5.**
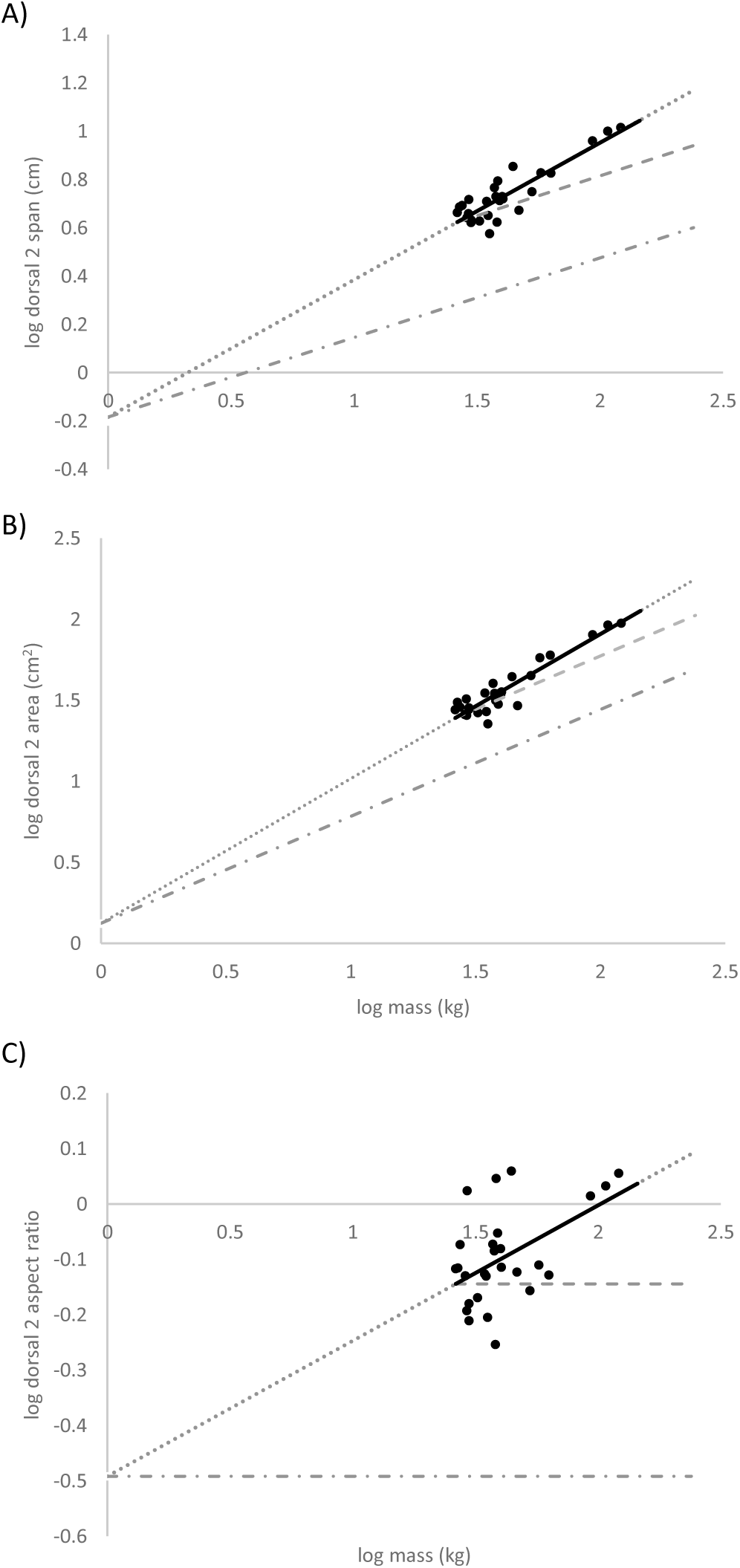
Allometric scaling of the second dorsal fin, represented by the black dots and solid line. Dotted line represents the allometric scaling extended across the size range observed in blue sharks (1kg-240kg). Dashed line represents isometric scaling initiated from the smallest shark observed in the dataset of this study (26kg). Dashed and dotted line represents isometric scaling initiated from the size of a blue shark pup (1 kg). A) Fin span (allometric slope = 0.57, R^2^ = 0.78; isometric slope = 0.33). B) Fin area (allometric slope = 0.89, R^2^ = 0.87; isometric slope = 0.66). C) Fin aspect ratio (allometric slope = 0.25, R^2^ = 0.23; isometric slope = 0).

**Figure 6.**
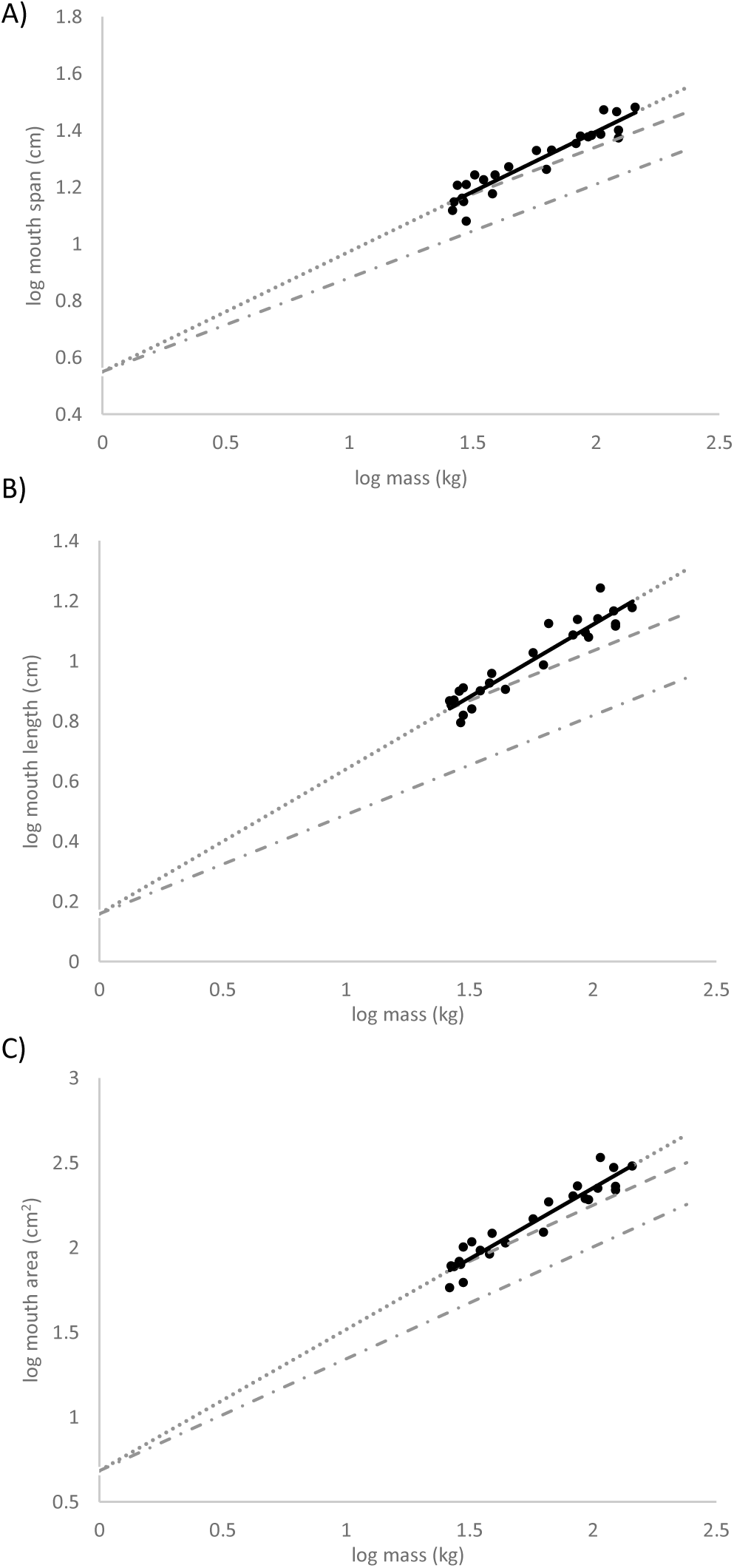
Allometric scaling of the mouth, represented by the black dots and solid line. Dotted line represents the allometric scaling extended across the size range observed in blue sharks (1kg-240kg). Dashed line represents isometric scaling initiated from the smallest shark observed in the dataset of this study (26kg). Dashed and dotted line represents isometric scaling initiated from the size of a blue shark pup (1 kg). A) Mouth span (allometric slope = 0.42, R^2^ = 0.88; isometric slope = 0.33). B) Mouth length (allometric slope = 0.48, R^2^ = 0.88; isometric slope = 0.33). C) Mouth area (allometric slope = 0.83, R^2^ = 0.88; isometric slope = 0.66).

**Table 1.** Scaling relationships from linear regressions for log10 transformed morphological variables vs. log10 transformed body mass (M).

| Variable | N | Range | Intercept | Slope ± CI | SE<sub>slope</sub> | Adj R<sup>2</sup> | P value | Scaling |
| --- | --- | --- | --- | --- | --- | --- | --- | --- |
| Standard Length | 36 | 140-278cm | 1.65 | 0.364 ± 0.0530 | 0.0261 | 0.915 | <0.001 | Isometry (M<sup>1/3</sup>) |
| Body Fineness Ratio | 36 | 5.95-8.15 | 0.757 | 0.0443 ± 0.0645 | 0.0318 | 0.0730 | 0.0608 | Isometry (M<sup>0</sup>) |
| Body Frontal Area | 36 | 270-959cm<sup>2</sup> | 1.51 | 0.690 ± 0.0882 | 0.0434 | 0.933 | <0.001 | Isometry (M<sup>2/3</sup>) |
| Body Surface Area | 26 | 6186-21626cm<sup>2</sup> | 2.64 | 0.796 ± 0.0481 | 0.0233 | 0.984 | <0.001 | Positive Allometry (M<sup>>2/3</sup>) |
| Body Volume | 26 | 19600-122000cm<sup>3</sup> | 2.58 | 1.18 ± 0.0680 | 0.0335 | 0.983 | <0.001 | Positive Allometry (M<sup>>1</sup>) |
| Body Density | 26 | 1040-1580kg·m<sup>-3</sup> | 3.42 | -0.181 ± 0.0690 | 0.0335 | 0.610 | <0.001 | Negative Allometry (M<sup><0</sup>) |
| Pitch & Yaw Inertia | 36 | 54.2-1130kg·m<sup>2</sup> | -0.697 | 1.72 ± 0.106 | 0.0522 | 0.983 | <0.001 | Isometry (M<sup>5/3</sup>) |
| Body Roll Inertia | 26 | 0.118-2.03kg·m<sup>2</sup> | -3.47 | 1.76 ± 0.0587 | 0.0285 | 0.994 | <0.001 | Positive Allometry (M<sup>>5/3</sup>) |
| Caudal Dorsal Span | 36 | 43.0-74.0cm | 1.23 | 0.292 ± 0.0682 | 0.0335 | 0.806 | <0.001 | Isometry (M<sup>1/3</sup>) |
| Caudal Dorsal Area | 36 | 238-975cm<sup>2</sup> | 1.62 | 0.629 ± 0.132 | 0.0649 | 0.837 | <0.001 | Isometry (M<sup>2/3</sup>) |
| Caudal Dorsal Aspect Ratio | 36 | 4.01-8.86 | 0.851 | -0.0439 ± 0.118 | 0.0580 | 0.00303 | 0.300 | Isometry (M<sup>0</sup>) |
| Caudal Ventral Span | 35 | 14.4-37.3cm | 0.574 | 0.458 ± 0.0837 | 0.0411 | 0.874 | <0.001 | Positive Allometry (M<sup>>1/3</sup>) |
| Caudal Ventral Area | 35 | 96.9-400cm<sup>2</sup> | 1.05 | 0.728 ± 0.157 | 0.0770 | 0.834 | <0.001 | Isometry (M<sup>2/3</sup>) |
| Caudal Ventral Aspect Ratio | 35 | 1.80-3.68 | 0.0961 | 0.188 ± 0.0980 | 0.0482 | 0.453 | <0.001 | Positive Allometry (M<sup>>0</sup>) |
| Dorsal 1 Span | 34 | 11.1-29.7cm | 0.244 | 0.581 ± 0.0842 | 0.0414 | 0.916 | <0.001 | Positive Allometry (M<sup>>1/3</sup>) |
| Dorsal 1 Area | 34 | 110-510cm<sup>2</sup> | 0.689 | 0.958 ± 0.134 | 0.0656 | 0.922 | <0.001 | Positive Allometry (M<sup>>2/3</sup>) |
| Dorsal 1 Aspect Ratio | 34 | 1.08-1.83 | -0.200 | 0.205 ± 0.0923 | 0.0453 | 0.5253 | <0.001 | Positive Allometry (M<sup>>0</sup>) |
| Dorsal 2 Span | 27 | 3.75-10.3cm | -0.184 | 0.568 ± 0.112 | 0.0546 | 0.775 | <0.001 | Positive Allometry (M<sup>>1/3</sup>) |
| Dorsal 2 Area | 27 | 22.5-94.0cm<sup>2</sup> | 0.124 | 0.892 ± 0.130 | 0.0626 | 0.866 | <0.001 | Positive Allometry (M<sup>>2/3</sup>) |
| Dorsal 2 Aspect Ratio | 27 | 0.556-1.14 | -0.492 | 0.245 ± 0.157 | 0.0762 | 0.226 | 0.00701 | Positive Allometry (M<sup>>0</sup>) |
| Pectoral Span | 34 | 33.7-63.4cm | 1.11 | 0.314 ± 0.0545 | 0.0267 | 0.887 | <0.001 | Isometry (M<sup>1/3</sup>) |
| Pectoral Area | 34 | 306-859cm<sup>2</sup> | 1.66 | 0.594 ± 0.112 | 0.0553 | 0.867 | <0.001 | Isometry (M<sup>2/3</sup>) |
| Pectoral Aspect Ratio | 34 | 3.59-5.24 | 0.564 | 0.0339 ± 0.0629 | 0.0309 | 0.0370 | 0.142 | Isometry (M<sup>0</sup>) |
| Head Span | 36 | 21.4-39.8cm | 0.899 | 0.319 ± 0.0603 | 0.0297 | 0.866 | <0.001 | Isometry (M<sup>1/3</sup>) |
| Head Area | 36 | 571-2230cm<sup>2</sup> | 1.73 | 0.731 ± 0.130 | 0.0642 | 0.877 | <0.001 | Isometry (M<sup>2/3</sup>) |
| Head Aspect Ratio | 36 | 0.618-1.42 | 0.0661 | -0.0962 ± 0.159 | 0.0781 | 0.0523 | 0.0960 | Isometry (M<sup>0</sup>) |
| Mouth Span | 25 | 12.0-30.2cm | 0.550 | 0.422 ± 0.080 | 0.0387 | 0.880 | <0.001 | Positive Allometry (M<sup>>1/3</sup>) |
| Mouth Length | 25 | 6.21-17.4cm | 0.158 | 0.481 ± 0.0926 | 0.0448 | 0.876 | <0.001 | Positive Allometry (M<sup>>1/3</sup>) |
| Mouth Area | 25 | 62.1-338cm<sup>2</sup> | 0.685 | 0.833 ± 0.105 | 0.143 | 0.907 | <0.001 | Positive Allometry (M<sup>>2/3</sup>) |
| Mouth Aspect Ratio | 25 | 2.33-3.34 | 0.417 | 0.0113 ± 0.0879 | 0.0433 | -0.0384 | 0.741 | Isometry (M<sup>0</sup>) |

**Table 2.** Results from two sample t-tests used to compare differences in the means for the variables related to swimming in larger versus smaller sharks, grouped in half based on the mass of the sharks.

| Variable | Scaling | Small Sharks (Mean ± SEM) | Large Sharks (Mean ± SEM) | p-value |
| --- | --- | --- | --- | --- |
| Body Frontal Area | Isometry | 11.1 ± 0.390 | 8.34 ± 0.220 | <0.001 |
| Body Surface Area | Allometry | 218 ± 3.64 | 179 ± 3.68 | <0.001 |
| Body Roll Inertia | Isometry | 0.00503 ± 0.000217 | 0.0106 ± 0.000901 | <0.001 |
| Body Pitch Inertia | Isometry | 2.588 ± 0.0759 | 4.952 ± 0.404 | <0.001 |
| Caudal Dorsal Span | Allometry | 1.51 ± 0.0401 | 0.805 ± 0.0520 | <0.001 |
| Caudal Dorsal Area | Isometry | 11.9 ± 0.429 | 8.10 ± 0.306 | <0.001 |
| Caudal Ventral Span | Allometry | 0.580 ± 0.0161 | 0.367 ± 0.0219 | <0.001 |
| Caudal Ventral Area | Isometry | 4.51 ± 0.192 | 3.506 ± 0.164 | <0.001 |
| Dorsal 1 Span | Allometry | 0.416 ± 0.0116 | 0.281 ± 0.0111 | <0.001 |
| Dorsal 1 Area | Allometry | 4.27 ± 0.156 | 4.11 ± 0.146 | 0.474 |
| Dorsal 2 Span | Allometry | 0.147 ± 0.00550 | 0.112 ± 0.00717 | 0.00103 |
| Dorsal 2 Area | Allometry | 0.921 ± 0.0330 | 0.865 ± 0.035 | 0.253 |
| Pectoral Span | Isometry | 1.21 ± 0.0379 | 0.665 ± 0.0439 | <0.001 |
| Pectoral Area | Isometry | 11.6 ± 0.350 | 7.66 ± 0.301 | <0.001 |

**Table 3.** Extent to which variables that scaled with allometry were larger or smaller than what would occur if they scaled with isometry (0% reference). Column 145kg*compares the largest shark in our dataset (145kg) relative to what would occur with isometric growth initiated from the smallest shark in our dataset (26kg) (see Figure 2-6). The other three columns display effects of allometric growth for the smallest shark in our dataset (26kg), the largest shark in our dataset (145kg), and the largest blue shark recorded in the World Record Game Fishes (240kg) (International Game Fish Association, 2002), relative to what would occur with isometric growth initiated from the mass of a typical-sized blue shark pup (∼1kg) (Kohler et al., 1992).

| Variable | 145kg* | 26kg | 145kg | 240kg |
| --- | --- | --- | --- | --- |
| Dorsal 1 Span | 35% | 56% | 71% | 75% |
| Dorsal 1 Area | 39% | 62% | 77% | 81% |
| Dorsal 1 Aspect Ratio | 30% | 49% | 64% | 67% |
| Dorsal 2 Span | 33% | 54% | 70% | 73% |
| Dorsal 2 Area | 33% | 53% | 69% | 72% |
| Dorsal 2 Aspect Ratio | 34% | 55% | 70% | 74% |
| Caudal Ventral Span | 20% | 34% | 47% | 51% |
| Caudal Ventral Aspect Ratio | 28% | 46% | 61% | 64% |
| Body Surface Area | 21% | 36% | 49% | 53% |
| Body Volume | 27% | 45% | 59% | 63% |
| Body Density | -27% | -45% | -59% | -63% |
| Body Roll Inertia | 16% | 28% | 40% | 43% |
| Mouth Span | 15% | 26% | 37% | 40% |
| Mouth Length | 23% | 39% | 53% | 56% |
| Mouth Area | 26% | 43% | 58% | 61% |

## DISCUSSION

Blue sharks demonstrate an extensive size range, from about 35-380 cm in length and 1kg-240kg in mass (Ebert et al., 2021; International Game Fish Association, 2002; Kohler et al., 1992). When morphology scales in an isometric fashion relative to increasing mass (M), linear (M^1/3^) and area (M^2/3^) measures decrease, while the turning inertia (M^5/3^) of the shark will increase, relative to mass. Hence, unless allometry changes these scaling relationships to promote a one-to-one relationship with mass, shifts in function in variables of morphology related to swimming performance and energetics are expected to ensue. The following discusses the functional implications of scaling relationships in various aspects of body and fin morphology relative to increasing mass in blue sharks.

### Body

As blue sharks grow, standard length increased with isometry, while body volume increased with positive allometry (Figure 1, Table 1). This was associated with an isometric increase in the fineness ratio, frontal area, and yaw and pitch inertia of the body (Table 1), whereas roll inertia and surface area of the body increased with positive allometry (Figure 2). The latter suggests that larger blue sharks require more deflecting force to roll, and generate more skin drag in swimming, compared to isometric scaling (Blake, 1983; Videler, 1993; Webb, 1975; Webb, 2005; Webb and Weihs, 2015). However, surface and frontal areas of the body were significantly less relative to mass in larger versus smaller sharks (Table 2), and consequently, the body form of larger blue sharks is expected to generate less pressure and skin (i.e. total) drag relative to mass than smaller sharks (Blake, 1983; Videler, 1993; Webb, 1975). Furthermore, yaw, pitch and roll inertia were significantly greater relative to mass in larger versus smaller blue sharks (Table 2). Hence, larger blue sharks are anticipated to require more force to deflect the body relative to mass, and therefore are expected to be more inherently stable than smaller blue sharks (Webb, 2005; Webb and Weihs, 2015). Increased inherent stability and decreased drag relative to body mass are likely to reduce the cost of transport in larger blue sharks in sustained rectilinear swimming, such as in long-distance migratory behaviours (Blake, 1983; Videler, 1993; Webb, 1975; Webb, 2005; Webb and Weihs, 2015).

Sharks are negatively buoyant, and the present results suggest the relative relationship between hydrostatic and hydrodynamic lift required to establish vertical equilibrium changes in blue sharks as they grow. The density of the body in blue sharks increased with negative allometry (Figure 2, Table 1). This corresponds with positive allometry in liver size of blue sharks with increasing mass in previous research (M^1.363^ ± 0.205 95% confidence interval) (Bone and Roberts, 1969), and that density of smaller blue sharks was found to be greater than larger blue sharks (Bone and Roberts, 1969). Therefore, larger blue sharks appear to demonstrate a shift towards neutral buoyancy, as predicted for other species (Bone and Roberts, 1969; Gleiss et al., 2017; Iosilevskii and Papastamatiou, 2016). These changes in density, and thus, greater relative contribution of hydrostatic lift in larger blue sharks, coincide with the allometric increase in span of the ventral lobe of the caudal fin discussed below (Figure 3, Table 1). Hence, hydrodynamic lift generated from the more symmetrical caudal fin is expected to be reduced as blue sharks grow (see Caudal Fin section below), but perhaps in turn is compensated for by the increase in hydrostatic lift. Shifting towards neutral buoyancy (i.e. increased hydrostatic lift) in larger sharks has been modeled to be associated with a reduced cost of transport for sustained swimming (Gleiss et al., 2017). Furthermore, a greater reliance on hydrodynamic vs hydrostatic lift is also likely to enhance burst maneuvering in small versus larger sharks, such as in escape responses, whereby negative buoyancy has been modeled to be more energetically efficient in accelerated movements (Gleiss et al., 2017). This corresponds with the expectation that smaller sharks are probably susceptible to relatively higher predation pressures than larger sharks (Chase, 1999; Guttridge et al., 2012; Heithaus, 2007; Heupel et al., 2007; Karythis et al., 2020), and therefore, may benefit from enhanced maneuverability despite the constraints of possible higher costs of transport.

### Caudal fin

The caudal fin generates primary thrust in swimming sharks, (Alexander, 1965; Ferry and Lauder, 1996; Flammang et al., 2011; Wilga and Lauder, 2002). Isometric growth was revealed in all aspects of the dorsal measurements of the caudal fin in blue sharks, whereas isometry was demonstrated in the ventral area, while the ventral span and aspect ratio of the caudal fin increased with positive allometry (Figure 3, Table 1). Relative to mass, larger blue sharks have significantly smaller spans and areas of the ventral and dorsal caudal fins compared to smaller blue sharks (Table 2). Furthermore, the positive allometry in the span of the ventral caudal fin shifts the shape of the entire caudal fin from heterocercal (i.e. asymmetrical) to more symmetrical in larger blue sharks. A heterocercal tail, common to most sharks, provides hydrodynamic lift by displacing water both backwards and downwards, whereby the reactive forces from the downwards vector of water shed from the tail lifts the body upwards in addition to generating thrust (Affleck, 1950; Alexander, 1965; Flammang et al., 2011; Lauder and Di Santo, 2015; Wilga and Lauder, 2002). In contrast, the more symmetrical caudal fin observed in larger blue sharks is anticipated to displace water backwards relative to forward movement and to a lesser extent downwards (Affleck, 1950; Flammang et al., 2011; Wilga and

Lauder, 2002). As discussed above, this is associated with negative allometric changes in the density of blue sharks as they grow, where they also become less negatively buoyant, and hence, may experience apparent shifts from the requirements for hydrodynamic toward more hydrostatic lift (see Density section above). A shift from a heterocercal tail to a symmetrical tail with increased body size has also been noted in several other species of sharks (Ahnelt et al., 2020; Fu et al., 2016; Gayford et al., 2023a; Irschick and Hammerschlag, 2015; Reiss and Bonnan, 2010; Sternes and Higham, 2022). Furthermore, an allometric increase in aspect ratio of the ventral lobe was also measured in blue sharks (Figure 3, Table 1), which is associated with increased hydrodynamic efficiency in the generation of thrust (Blake, 1983; Videler, 1993; Webb, 1975). Overall, a more symmetrical caudal fin, an increase in aspect ratio of the ventral fin, and smaller fins relative to mass are features associated with the capacity for higher sustained swimming speeds and lower costs of transport in migration (Blake, 1983; Iliou et al., 2023; Maia et al., 2012; Videler, 1993; Webb, 1975; Wilga and Lauder, 2002). In contrast, sharks with greater asymmetry in the caudal fin, a smaller aspect ratio in the ventral fin, and larger fins relative to mass appear to be associated with the capacity for slower sustained swimming speeds, but higher maneuverability (Alexander, 1965; Blake, 1983; Iliou et al., 2023; Maia et al., 2012; Videler, 1993; Webb, 1975; Wilga and Lauder, 2002). Scaling of the caudal fin that reduces the cost of transport of migration, even at the cost of reduced maneuverability, is likely beneficial for larger blue sharks that probably experience less predation pressure (Chase, 1999; Guttridge et al., 2012; Heithaus, 2007; Heupel et al., 2007; Karythis et al., 2020).

### Dorsal fins

The dorsal fins predominately act as stabilizers, particularly in roll rotation, and also potentially generate secondary thrust (Lingham-soliar, 2005; Maia and Wilga, 2013b; Maia and Wilga, 2013a; Standen, 2005; Standen and Lauder, 2007). Positive allometry was measured in the span, area and aspect ratio of both the anterior and posterior dorsal fins (Figure 4 and 5, Table 1). While there was a significant difference in span, there was not a significant difference in area of the dorsal fins relative to mass in larger versus smaller blue sharks (Table 2).

Furthermore, the positive allometry in the scaling of span would provide a disproportionally greater moment arm to promote greater roll stability of the body in larger blue sharks, compared to isometric growth. Positive allometry in the scaling of area is expected to promote a disproportionate increase in the capacity for the generation of torque and thrust from these fins in larger blue sharks, compared to isometric growth. Hence, allometric changes in the dorsal fins might act to maintain the function of these fins relative to mass, particularly in force generation for roll stability. These allometric changes in both the span and area of the dorsal fins are anticipated to be especially beneficial in potentially providing larger blue sharks with enhanced stabilization of the body when swimming at high sustained speeds because of the shift in aspect ratio (Blake, 1983; Lighthill, 1969; Webb, 1975; Webb, 1984). A positive allometric increase in the aspect ratio (span^2^/area) of the dorsal fins might bestow larger blue sharks with the capacity to migrate with a lower cost of transport and achieve higher sustained swimming speeds because they generate less drag for a given amount of lift (Blake, 1983; Lighthill, 1969; Webb, 1975). However, with less capacity to generate drag, fins with higher aspect ratios are considered less favourable in the performance of unsteady burst swimming such as in escape behaviours from larger predators (Blake, 1983; Domenici et al., 2004; Seamone et al., 2014; Walker and Westneat, 2000; Webb, 1975; Webb, 1984).

### Pectoral fins

Pectoral fins are important in promoting turning maneuvers in sharks (Drucker, 2003; Drucker and Lauder, 2001; Hoffmann et al., 2019). All aspects of the pectoral fins in blue sharks scaled with isometry (Table 1), and furthermore, the span and area of these fins was significantly smaller in larger versus smaller blue sharks relative to mass (Table 2). While shape of the pectoral fins remains proportional with growth, smaller fins relative to mass probably benefit blue sharks in reducing drag and hence the cost of transport in long-distance rectilinear migrations, whereas larger fins relative to mass likely benefit smaller blue sharks in maneuvering, such as in fast start escape from larger predators (Blake, 1983; Videler, 1993; Webb, 1975; Webb, 2005; Webb and Weihs, 2015).

It has been suggested that changes in habitat that impose different requirements for maneuvering with growth might promote allometric scaling relationships in the aspect ratio of the pectoral fins in sharks (Sternes and Higham, 2022). Hammerhead sharks demonstrate profound shifts in habitat as they grow, migrating from shallow coastal habitats to deep bodied oceanic waters beyond the continental shelf (Clarke, 1971; Colombo Estupiñán-Montaño et al., 2021; Duncan and Holland, 2006; Hoyos-Padilla et al., 2014). This change in habitat may underly a major difference between blue sharks and hammerheads in the scaling of the aspect ratio of pectoral fins with growth; hammerhead sharks showed positive allometry (Sternes and Higham, 2022), while blue sharks showed isometry. Shifting from shallow mangrove to oceanic habitats is expected to present different physical constraints and selective pressures on turning performance throughout the life history in hammerheads (Sternes and Higham, 2022). Pectoral fins with a lower aspect ratio observed in smaller hammerheads are more effective for generating torque and therefore maneuvering through dense mangrove environments, whereas high aspect ratio fins observed in larger hammerheads are energetically more economic for sustained migrations through wide-open oceanic environments (Blake, 1983; Videler, 1993; Webb, 1975). In blue sharks, the minimal physical constraints imposed by an oceanic environment remain relatively constant throughout their life history. Thus, there may not be strong selective pressures for allometric changes in pectoral fins with growth in blue sharks, as there appears to be in hammerheads (Iosilevskii and Papastamatiou, 2016).

### Head and mouth

The head and mouth are some of the most defining features of sharks, enabling them to effectively grasp prey and achieve some of the highest tropic levels of organisms in the ocean (Cullen and Marshall, 2019; Martin et al., 2005; Moyer and Dodd, 2023; Wilga and Motta, 2000). Allometric changes in the head and mouth with growth have been revealed in several shark species, which are expected to have impacts on predation strategies and ability to consume prey throughout their life history (Ahnelt et al., 2020; Fu et al., 2016; Gayford et al., 2023a; Sternes and Higham, 2022; Yun and Watanabe, 2023). For example, the heads of juvenile tiger sharks are more conical, which transition to relatively broader heads over ontogeny (Fu et al., 2016). This has been attributed to adult tiger sharks expanding their diet to consume larger and more diverse prey with age (e.g., turtles, mammals, and elasmobranchs), which requires greater bite area and force to process (Fu et al., 2016). In contrast, blue sharks are apparently generalist predators throughout their life history (Bazzi et al., 2021; Henderson et al., 2001; McCord and Campana, 2003; Moyer and Dodd, 2023; Young et al., 2010). The trophic level of blue sharks has been demonstrated to increase in some populations, which has been ascribed to variation in prey abundance in different feeding grounds and the ability for larger blue sharks to predate on larger prey (Hernandez-Aguilar et al., 2016; Rabehagasoa et al., 2012). While the basic morphology of the head of blue sharks scaled with isometry, the width, length and area of the mouth scaled with positive allometry (Figure 6, Table 1). Hence, larger blue sharks are anticipated to have an ability to grasp and consume disproportionately larger prey, compared to if the mouth scaled isometrically. This might be beneficial for increasing predatory success in opportunistic prey availability.

### Relative change in allometric growth versus isometric growth

The relative change in variables that scaled with allometry in blue sharks, as opposed to scaling with isometry, depends on the point that allometric growth was initiated in development, in addition to the size of the shark in which comparisons were made (Figure 2-6, Table 3). The magnitude of change was greatest in scenarios where allometry was initiated from the size of a pup (1kg) and compared with measures in the largest blue shark recorded in the population (240kg) (International Game Fish Association, 2002). Relative changes in this scenario were substantial for all variables that scaled allometrically in blue sharks, ranging from 40-80% compared with isometry (Table 3). That said, it is highly likely that allometric growth might be initiated or terminated at a later or earlier point in the development of the shark, respectively. For example, when the allometric curve for density was extended across the full-size range of the blue shark population, it provided unreasonable measures of density, whereby blue shark pups were likely too dense, and sharks larger than 180kg were predicted to become less dense than seawater (Figure 2); hence, changes in body density around 63% in blue sharks is unlikely. Therefore, future allometry studies that encompass the entire size range of an organism and include large sample sizes may consider exploring scaling relationships within different ontogenetic stages. This is expected to be promising in determining when in development allometric scaling is initiated, and furthermore, in generating accurate measures of the relative changes in variables that scale allometrically. Nevertheless, when the extent of the allometric changes were considered solely within the size range of the dataset from this study, which is expected to be an underestimate, noteworthy changes of 15-40% were still revealed (Figure 2-6, Table 3). The magnitude of change for allometric parameters in blue sharks probably fall somewhere in between these two scenarios. Thus, allometric changes such as in the shape of the fins and body density in blue sharks are predicted to have substantial impacts on the biology of blue sharks compared to if these features scaled isometrically.

### Summary and conclusions

Selective pressures related to ontogenetic shifts in habitat and diet have been proposed to promote allometric changes in morphology with growth in sharks (Fu et al., 2016; Gayford et al., 2023a; Lingham-Soliar, 2005; Sternes and Higham, 2022). Blue sharks are generalist predators that never cease swimming in a oceanic environment throughout their life history (Bazzi et al., 2021; Henderson et al., 2001; McCord and Campana, 2003; Moyer and Dodd, 2023; Queiroz et al., 2005; Stevens, 1976; Vandeperre et al., 2014; Young et al., 2010). Hence, we hypothesized that the form of the body and shape of the fins would scale with isometry with increasing size in blue sharks because they do not appear to experience major changes in the physical constraints imposed by environment or prey type. Although blue sharks demonstrated isometry in many aspects of the body and fins, our hypothesis was rejected based on several measures of allometric changes (Figure 2-6, Table 1), supporting the prospect of a niche shift in blue sharks (Gayford et al., 2023a). Indeed, larger blue sharks increase their migratory range in the Atlantic Ocean compared to smaller individuals, and stable isotope studies in blue sharks demonstrate differences between populations, whereby trophic level and feeding grounds may vary with size despite the lack of profound changes in diet throughout their life history (Hernandez-Aguilar et al., 2016; Kiszka et al., 2015; Li et al., 2016; Macneil et al., 2005; Rabehagasoa et al., 2012; Vidal et al., 2023).

We also demonstrated that through isometry, larger blue sharks grow into significantly smaller fins, bodies with less area, and bodies that are inherently more stable relative to mass (Table 2). Hence, isometric scaling does not necessarily imply functional similarity, and allometric scaling may sometimes be critical in actually conserving, rather than altering, function relative to mass. Both allometric and isometric results in this study suggest that larger individuals might have the capacity to achieve higher sustained speeds and lower costs of transport in migration, particularly due to the predictions of decreased drag in the body and fins, and bodies that are more inherently stable, but that smaller blue sharks may be more adept in unsteady swimming such as escape responses (Domenici, 2001; Gleiss et al., 2017; Webb, 1975). Similar predictions for the constraints of performance in unsteady maneuverability with increasing size, due to the relative scaling of control surface area and body mass, have been made for rorqual whales (Kahane-Rapport and Goldbogen, 2018). Blue sharks face predation pressures from larger predators, such as marine mammals and sharks (Ford et al., 2011; McCord and Campana, 2003). Given the exceptional size range of blue sharks (35-380cm in length and 1-240kg in mass), we predict that changes with growth that increase the capacity for economic sustained swimming, even at the functional cost of reduced maneuverability, may be a potential advantage in larger blue sharks if predation pressures are reduced.

## ACKNOWLEDGMENTS

This research was supported by a Natural Sciences and Engineering Research Council of Canada Discovery Grant to DAS and a NSERC Alexander Graham Bell and University of Calgary Killam Scholarship to SGS. We would like to thank the Department of Fisheries and Oceans, Canada, and the organizers of the Petit-de-Grat, Lockeport and Louisbourg shark derbies for allowing us to collect data. Further, we would like to thank R.G. Argand, and A.J. Laughlin, C.A. Seamone, and C.D. Seamone for their substantial help with collecting data at these derbies.

## REFERENCES

Affleck (1950). Some points in the function, development and evolution of the tail in fishes. In Proceedings of the zoological society of London, pp. 349–368. Oxford, UK: Blackwell Publishing Ltd.

Ahnelt, H., Sauberer, M., Ramler, D., Koch, L. and Pogoreutz, C. (2020). Negative allometric growth during ontogeny in the large pelagic filter-feeding basking shark. Zoomorphology 139, 71–83.

Alexander, R. M. (1965). The lift produced by the heterocercal tails of Selachii. J. Exp. Biol. 43, 131–138.

Bazzi, M., Campione, N. E., Kear, B. P., Pimiento, C. and Ahlberg, P. E. (2021). Feeding ecology has shaped the evolution of modern sharks. Curr. Biol. 31, 5138–5148.e4.

Bellodi, A., Mulas, A., Daniel, L., Cau, A., Porcu, C., Carbonara, P. and Follesa, M. C. (2023). Ontogenetic shifts in body morphology of demersal sharks’ species (order: Squaliformes) inhabiting the western-central Mediterranean Sea, with implications for their bio-ecological role. Biology (Basel). 12, 12.

Birn-Jeffery, A. V. and Higham, T. E. (2014). The scaling of uphill and downhill locomotion in legged animals. Integr. Comp. Biol. 54, 1159–1172.

Blake, R. W. (1983). Fish locomotion. Cambridge, UK: Cambridge University Press.

Bone, Q. and Roberts, B. L. (1969). The density of elasmobranchs. J. Mar. Biol. Assoc. United Kingdom 49, 913–937.

Campana, S. E., Dorey, A., Fowler, M., Joyce, W., Wang, Z., Wright, D. and Yashayaev, I. (2011). Migration pathways, behavioural thermoregulation and overwintering grounds of blue sharks in the Northwest Atlantic. PLoS One 6,.

Chase, J. M. (1999). Food web effects of prey size refugia: Variable interactions and alternative stable equilibria. Am. Nat. 154, 559–570.

Clarke, T. (1971). Ecology of scalloped hammerhead shark, Sphyrna-Lewini, in Hawaii. Pacific Sci. 25, 133-.

Colombo Estupiñán-Montaño, Galván-Magaña, F., Elorriaga-Verplancken, F., Zetina-Rejón, M. J. A. S.-G., Polo-Silva, C. J., Villalobos-Ramírez, D. J., Rojas-Cundumí, J. and Delgado-Huertas, A. (2021). Ontogenetic feeding ecology of the scalloped hammerhead shark *Sphyrna lewini* in the Colombian Eastern Tropical Pacific. Mar. Ecol. Prog. Ser. 663, 127– 143.

Cook, A. (1996). Ontogeny of feeding morphology and kinematics in juvenile fishes: a case study of the cottid fish *Clinocottus analis*. J. Exp. Biol. 199, 1961–71.

Cullen, J. A. and Marshall, C. D. (2019). Do sharks exhibit heterodonty by tooth position and over ontogeny ? A comparison using elliptic Fourier analysis. 687–700.

Di Santo, V., Kenaley, C. P. and Lauder, G. V. (2017). High postural costs and anaerobic metabolism during swimming support the hypothesis of a U-shaped metabolism–speed curve in fishes. Proc. Natl. Acad. Sci. U. S. A. 114, 13048–13053.

Dial, K. P., Greene, E. and Irschick, D. J. (2008). Allometry of behavior. Trends Ecol. Evol. 23, 394–401.

Domenici, P. (2001). The scaling of locomotor performance in predator-prey encounters: from fish to killer whales. Comp. Biochem. Physiol. - A Mol. Integr. Physiol. 131, 169–182.

Domenici, P. and Blake, R. W. (1997). The kinematics and performance of fish fast-start swimming. J. Exp. Biol. 200, 1165–1178.

Domenici, P., Standen, E. M. M. and Levine, R. P. (2004). Escape manoeuvres in the spiny dogfish (*Squalus acanthias*). J. Exp. Biol. 207, 2339–2349.

Drucker, E. G. (2003). Function of pectoral fins in rainbow trout: behavioral repertoire and hydrodynamic forces. J. Exp. Biol. 206, 813–826.

Drucker, E. G. and Lauder, G. V. (2001). Wake dynamics and fluid forces of turning maneuvers in sunfish. J. Exp. Biol. 204, 431–442.

Duncan, K. M. and Holland, K. N. (2006). Habitat use, growth rates and dispersal patterns of juvenile scalloped hammerhead sharks *Sphyrna lewini* in a nursery habitat. Mar. Ecol. Prog. Ser. 312, 211–221.

Ebert, D., Dando, M. and Fowler, S. (2021). Sharks of the world: a complete guide. Princeton: Princeton University Press.

Enquist, B. J., Economo, E. P., Huxman, T. E., Allen, A. P., Ignace, D. D. and Gillooly, J. F. (2003). Scaling metabolism from organisms to ecosystems. Nature 423, 639–642.

Estrada, J. A., Rice, A. N., Lutcavage, M. E. and Skomal, G. B. (2003). Predicting trophic position in sharks of the north-west Atlantic Ocean using stable isotope analysis. J. Mar. Biol. Assoc. UK 83, 1347–1350.

Ferry, L. and Lauder, G. V. (1996). Heterocercal tail function in leopard sharks: a three-dimensional kinematic analysis of two models. J. Exp. Biol. 199, 2253–68.

Flammang, B. E., Lauder, G. V., Troolin, D. R. and Strand, T. (2011). Volumetric imaging of shark tail hydrodynamics reveals a three-dimensional dual-ring vortex wake structure. Proc. R. Soc. B Biol. Sci. 278, 3670–3678.

Ford, J. K. B., Ellis, G. M., Matkin, C. O., Wetklo, M. H., Barrett-Lennard, L. G. and Withler, R. E. (2011). Shark predation and tooth wear in a population of northeastern pacific killer whales. *Aquat*. Biol. 11, 213–224.

Fu, A. L., Hammerschlag, N., Lauder, G. V., Wilga, C. D., Kuo, C. Y. and Irschick, D. J. (2016). Ontogeny of head and caudal fin shape of an apex marine predator: the tiger shark (*Galeocerdo cuvier*). J. Morphol. 277, 556–564.

Galileo, G. (1914). 1637 Dialogues Concerning Two New Sciences. Translated by Henry Crew and A. DeSalvio. New York: The Macmillan Company.

Gamperl, A. K., Rodnick, K. J., Faust, H. A., Venn, E. C., Bennett, M. T., Crawshaw, L. I., Keeley, E. R., Powell, M. S. and Li, H. W. (2002). Metabolism, swimming performance, and tissue biochemistry of high desert redband trout (*Oncorhynchus mykiss ssp.*): evidence for phenotypic differences in physiological function. Physiol. Biochem. Zool. 75, 413–431.

Gayford, J. H., Whitehead, D. A., Ketchum, J. T. and Field, D. J. (2023a). The selective drivers of allometry in sharks (Chondrichthyes: Elasmobranchii). Zool. J. Linn. Soc. 1–21.

Gayford, J. H., Godfrey, H. and Whitehead, D. A. (2023b). Ontogenetic morphometry of the brown smoothhound shark *Mustelus henlei* with implications for ecology and evolution. 1–12.

Gayon, J. (2000). History of the concept of allometry. Am. Zool. 40, 748–758.

Gleiss, A. C., Potvin, J. and Goldbogen, J. A. (2017). Physical trade-offs shape the evolution of buoyancy control in sharks. Proc. R. Soc. B Biol. Sci. 284, 20171345.

Gould, S. J. (1966). Allometry and size in ontogeny and phylogeny. Biol. Rev. 41, 587–638.

Guttridge, T. L., Gruber, S. H., Franks, B. R., Kessel, S. T., Gledhill, K. S., Uphill, J., Krause, J. and Sims, D. W. (2012). Deep danger: intra-specific predation risk influences habitat use and aggregation formation of juvenile lemon sharks *Negaprion brevirostris*. Mar. Ecol. Prog. Ser. 445, 279–291.

Heithaus, M. R. (2007). Nursery areas as essential shark habitats: a theoretical perspective. Am. Fish. Soc. Symp. 50, 3–13.

Henderson, A. C., Flannery, K. and Dunne, J. (2001). Observations on the biology and ecology of the blue shark in the North-east Atlantic. J. Fish Biol. 58, 1347–1358.

Hernandez-Aguilar, S. B., Escobar-Sanchez, O., Galvan-Magana, F. and Abitia-Cardenaz, L. A. (2016). Trophic ecology of the blue shark ( *Prionace glauca*) based on stable isotopes ( d 13 C and d 15 N) and stomach content. J. Mar. Biol. Assoc. United Kingdom 96, 1403– 1410.

Heupel, M. R., Carlson, J. K. and Simpfendorfer, C. A. (2007). Shark nursery areas: concepts, definition, characterization and assumptions. 337, 287–297.

Higham, T. E., Seamone, S. G., Arnold, A., Toews, D., Janmohamed, Z., Smith, S. J. and Rogers, S. M. (2018). The ontogenetic scaling of form and function in the spotted ratfish, Hydrolagus colliei (Chondrichthyes: Chimaeriformes): fins, muscles, and locomotion. J. Morphol. 279, 1408–1418.

Hoffmann, S. L. and Porter, M. E. (2019). Body and pectoral fin kinematics during routine yaw turning in bonnethead sharks (*Sphyrna tiburo*). Integr. Org. Biol. 1,.

Hoffmann, S. L., Donatelli, C. M., Leigh, S. C., Brainerd, E. L. and Porter, M. E. (2019). Three-dimensional movements of the pectoral fin during yaw turns in the Pacific spiny dogfish, *Squalus suckleyi*. Biol. Open 8,.

Hoyos-Padilla, E. M., Ketchum, J. T., Klimley, A. P. and Galván-Magaña, F. (2014). Ontogenetic migration of a female scalloped hammerhead shark *Sphyrna lewini* in the Gulf of California. *Anim*. Biotelemetry 2, 1–9.

Huxley, J. S. and Tessier, G. (1936). Terminology of relative growth. Nature 137, 780–781.

Iliou, A. S., Vanderwright, W., Harding, L., Jacoby, D. M. P., Payne, N. L. and Dulvy, N. K. (2023). Tail shape and the swimming speed of sharks.

International Game Fish Association (2002). World Record Game Fishes.

Iosilevskii, G. and Papastamatiou, Y. P. (2016). Relations between morphology, buoyancy and Relations between morphology, buoyancy and energetics of requiem sharks.

Irschick, D. J. and Hammerschlag, N. (2015). Morphological scaling of body form in four shark species differing in ecology and life history. Biol. J. Linn. Soc. 144, 126–135.

Irschick, D. J., Fu, A., Lauder, G. V., Wilga, C. D., Kuo, C. and Hammerschlag, N. (2017). A comparative morphological analysis of body and fin shape for eight shark species. Biol. J. Linn. Soc. 122, 1–16.

James, A. E., Aaron, N. R., Lisa, J. N. and Gregory, B. S. (2006). Use of isotopic analysis of vertebrae in reconstructing ontogenetic feeding ecology in white sharks. Ecology 87, 829– 834.

Kahane-Rapport, S. R. and Goldbogen, J. A. (2018). Allometric scaling of morphology and engulfment capacity in rorqual whales. J. Morphol. 279, 1256–1268.

Karythis, S., Cornwell, T. O., Noya, L. G., McCarthy, I. D., Whiteley, N. M. and Jenkins, S. R. (2020). Prey vulnerability and predation pressure shape predator-induced changes in O_2_ consumption and antipredator behaviour. Anim. Behav. 167, 13–22.

Killen, S. S., Costa, I., Brown, J. A. and Gamperl, A. K. (2007). Little left in the tank: metabolic scaling in marine teleosts and its implications for aerobic scope. Proc. R. Soc. B Biol. Sci. 274, 431–438.

Kiszka, J. J., Aubail, A., Hussey, N. E., Heithaus, M. R., Caurant, F. and Bustamante, P. (2015). Plasticity of trophic interactions among sharks from the oceanic south-western Indian Ocean revealed by stable isotope and mercury analyses. Deep. Res. Part I 96, 49–58.

Kohler, N. E., Casey, J. G. and Turner, P. A. (1992). Length-length and length-weight relationships for 13 shark species from the Western North Atlantic. NOAA Tech. Memo. NMFS-NE*-*110 22, 1–22.

Lauder, G. V and Di Santo, V. (2015). Swimming mechanics and energetics of elasmobranch fishes. In Fish Physiology: Physiology of Elamobranch Fishes (ed. Shadwick, R. E.), Farrell, A. P.), and Brauner, C. J.), pp. 219–249. Elsevier.

Li, Y., Zhang, Y. and Dai, X. (2016). Trophic interactions among pelagic sharks and large predatory teleosts in the northeast central Pacific. J. Exp. Mar. Bio. Ecol. 483, 97–103.

Lighthill, M. J. (1969). Hydromechanics of aquatic animal propulsion. Annu. Rev. Fluid Mech. 1, 413–446.

Lingham-soliar, T. (2005). Dorsal fin in the white shark, Carcharodon carcharias: a dynamic stabilizer for fast swimming. 11, 1–11.

Lingham-Soliar, T. (2005). Caudal fin allometry in the white shark *Carcharodon carcharias*: implications for locomotory performance and ecology. Naturwissenschaften 92, 231–236.

Lowe, C. G., Wetherbee, B. M., Crow, G. L. and Tester, A. (1996). Ontogenetic dietary shifts and feeding behavior of the tiger shark, *Galeocerdo cuvier*, in Hawaiian waters. Environ. Biol. Fishes 47, 203–211.

Macneil, M. A., Skomal, G. B. and Fisk, A. T. (2005). Stable isotopes from multiple tissues reveal diet switching in sharks. 302, 199–206.

Maia, A. and Wilga, C. D. (2013a). Function of dorsal fins in bamboo shark during steady swimming. Zoology 116, 224–231.

Maia, A. and Wilga, C. D. (2013b). Anatomy and muscle activity of the dorsal fins in bamboo sharks and spiny dogfish during turning maneuvers. J. Morphol. 274, 1288–1298.

Maia, A., Wilga, C. D. and Lauder, G. V. (2012). Biomechanics of locomotion in sharks, rays, and chimaeras. In Biology of sharks and their relatives, second edition (ed. Carrier, J.), Musick, J.), and Heithaus, M.), p. 125‒151. Boca Raton.

Martin, R. A., Hammerschlag, N., Collier, R. S. and Fallows, C. (2005). Predatory behaviour of white sharks (*Carcharodon carcharias*) at Seal Island, South Africa. J. Mar. Biol. Assoc. UK 85, 1121–1135.

McCord, M. E. and Campana, S. E. (2003). A quantitative assessment of the diet of the blue shark (*Prionace glauca*) off Nova Scotia, Canada. J. Northwest Atl. Fish. Sci. 32, 57–63.

Moyer, J. K. and Dodd, J. (2023). Feeding kinematics and ethology of blue sharks, Prionace glauca (Carcharhiniformes: Carcharhinidae). Environ. Biol. Fishes.

Norin, T. and Gamperl, A. K. (2018). Metabolic scaling of individuals vs. populations: evidence for variation in scaling exponents at different hierarchical levels. Funct. Ecol. 32, 379–388.

Preti, A., Soykan, C. U., Dewar, H., Wells, R. J. D., Spear, N. and Kohin, S. (2012). Comparative feeding ecology of shortfin mako, blue and thresher sharks in the California Current. Environ. Biol. Fishes 95, 127–146.

Queiroz, N., Lima, F. P., Maia, A., Ribeiro, P. A., Correia, J. P. and M. Santos, A. (2005). Movement of blue shark, Prionace glauca, in the north-east Atlantic based on mark-recapture data. J. Mar. Biol. Assoc. United Kingdom 85, 1107–1112.

Rabehagasoa, N., Lorrain, A., Bach, P., Potier, M., Jaquemet, S., Richard, P. and Ménard, F. (2012). Isotopic niches of the blue shark Prionace glauca and the silky shark *Carcharhinus falciformis* in the south-western Indian Ocean. Endanger. Species Res. 17, 83–92.

Reiss, K. L. and Bonnan, M. F. (2010). Ontogenetic scaling of caudal fin shape in Squalus acanthias (chondrichthyes, elasmobranchii): a geometric morphometric analysis with implications for caudal fin functional morphology. Anat. Rec. 293, 1184–1191.

Schmidt-Nielsen, K. (1975). Scaling in biology: the consequences of size. J. Exp. Zool. 194, 287– 307.

Seamone, S. G., Blaine, T. and Higham, T. E. (2014). Sharks modulate their escape behavior in response to predator size, speed and approach orientation. Zoology 117, 377–382.

Sepulveda, C. A., Graham, J. B. and Bernal, D. (2007). Aerobic metabolic rates of swimming juvenile mako sharks, *Isurus oxyrinchus*. Mar. Biol. 152, 1087–1094.

Standen, E. M. (2005). Dorsal and anal fin function in bluegill sunfish *Lepomis macrochirus*: three-dimensional kinematics during propulsion and maneuvering. J. Exp. Biol. 208, 2753– 2763.

Standen, E. M. (2008). Pelvic fin locomotor function in fishes: three-dimensional kinematics in rainbow trout (*Oncorhynchus mykiss*). J. Exp. Biol. 211, 2931–2942.

Standen, E. M. and Lauder, G. V. (2007). Hydrodynamic function of dorsal and anal fins in brook trout (*Salvelinus fontinalis*). J. Exp. Biol. 210, 325–339.

Sternes, P. C. and Higham, T. E. (2022). Hammer it out: shifts in habitat are associated with changes in fin and body shape in the scalloped hammerhead (*Sphyrna lewini*). Biol. J. Linn. Soc. 136, 201–212.

Stevens, J. D. (1976). First results of shark tagging in the north-east atlantic, 1972–1975. J. Mar. Biol. Assoc. United Kingdom 56, 929–937.

Vandeperre, F., Aires-da-Silva, A., Fontes, J., Santos, M., Serrão Santos, R. and Afonso, P. (2014). Movements of blue sharks (Prionace glauca) across their life history. PLoS One 9,.

Vidal, A., Cardador, L., Garcia-Barcelona, S., Macias, D., Druon, J. N., Coll, M. and Navarro, J. (2023). The relative importance of biological and environmental factors on the trophodynamics of a pelagic marine predator, the blue shark (*Prionace glauca*). Mar. Environ. Res. 183, 105808.

Videler, J. J. (1993). Fish Swimming. Fish and F. London: Chapman and Hall.

Vogel, S. (2005). Living in a physical world III. Getting up to speed. J. Biosci. 30, 303–312.

Walker, J. A. and Westneat, M. W. (2000). Mechanical performance of aquatic rowing and flying. Proc. R. Soc. B Biol. Sci. 267, 1875–1881.

Watanabe, Y. Y., Payne, N. L., Semmens, J. M., Fox, A. and Huveneers, C. (2019). Swimming strategies and energetics of endothermic white sharks during foraging. J. Exp. Biol. 222,.

Webb, P. W. (1975). Hydrodynamics and Energetics of Fish Propulsion. Bull. Fish. Res. Board Canada 1–158.

Webb, P. W. (1984). Body form, locomotion and foraging in aquatic vertebrates. Am. Zool. 24, 107–120.

Webb, P. W. (2005). Stability and maneuverability. In Fish Physiology: Fish Biomechanics, pp. 281–332.

Webb, P. W. and Weihs, D. (2015). Stability versus maneuvering: Challenges for stability during swimming by fishes. Integr. Comp. Biol. 55, 753–764.

Wilga, C. D. and Lauder, G. V. (2000). Three-dimensional kinematics and wake structure of the pectoral fins during locomotion in leopard sharks *Triakis semifasciata*. J. Exp. Biol. 203, 2261–78.

Wilga, C. D. and Lauder, G. V. (2002). Function of the heterocercal tail in sharks: quantitative wake dynamics during steady horizontal swimming and vertical maneuvering. J. Exp. Biol. 205, 2365–2374.

Wilga, C. D. and Motta, P. J. (2000). Durophagy in sharks: Feeding mechanics of the hammerhead *Sphyrna tiburo*. J. Exp. Biol. 203, 2781–2796.

Young, J. W., Lansdell, M. J., Campbell, R. A., Cooper, S. P., Juanes, F. and Guest, M. A. (2010). Feeding ecology and niche segregation in oceanic top predators off eastern Australia. Mar. Biol. 157, 2347–2368.

Yun, C. and Watanabe, Y. Y. (2023). Allometric growth of the enigmatic deep-sea megamouth shark *Megachasma pelagios* Taylor, Compagno, and Struksaker, 1983 (Lamniformes, Megachasmidae). Fishes 8,.

